# Multiscale characterization of the mechanics of curved fibered structures with application to biological materials

**DOI:** 10.1101/2024.01.09.574800

**Authors:** J.A. Sanz-Herrera, A. Apolinar-Fernandez, A. Jimenez-Aires, P. Perez-Alcantara, J. Dominguez, E. Reina-Romo

**Author notes:** Corresponding author. Camino de los descubrimientos s/n, 41092 Seville, Spain.; *Email address:* (J.A. Sanz-Herrera).

## Abstract

Curved fibered structures are ubiquitous in nature and this organization is found in the majority of biological tissues. Indeed, the mechanical behavior of these materials is of pivotal importance in biomechanics and mechanobiology fields. In this paper, we develop a multiscale formulation to characterize the macroscopic mechanical nonlinear behavior from the microstructure of fibered matrices. From the analysis of the mechanics of a randomly curved single fiber, a fibered matrix model is built to determine the macroscopic behavior following a homogenization approach. The model is tested for tensile, compression and shear loads in a number of applications reminiscent to collagen extracellular matrices. However, any other fibered microstructures can be studied following the proposed formulation. The presented approach naturally recovers instabilities at compression as well as the strain stiffening regime, which are observed experimentally in the mechanical behavior of collagen matrices. Indeed, it was found that the bending energy associated to fiber unrolling, is the most important source of energy developed by fibers for the analyzed cases in tensile and shear in all deformation regions (except the strain stiffening region), whereas bending energy dominates at compression too during buckling. The proposed computational framework can also be used to perform multiscale simulations in the referred applications. As a result, the developed methodology may be an interesting and complementary tool to characterize the nonlinear behavior and evolution of curved fibered structures present in biology and engineered materials.

## 1. Introduction

Fibrous networks play a fundamental role in many biological materials. Fibered tissues like the extracellular matrix (ECM) of biological tissues are composed by fibrils (e.g. collagen) connected to each other and surrounding nonfibrillar matrix. This hierarchical structure makes fibered materials macroscopic behavior to be largely dependent on its microstructural organization. Therefore, it is important to understand and characterize the behavior of fibrous tissues to provide insights into the pathophysiology of different diseases such as cancer or arteriosclerosis [1]. In many solid tumors there is a significant collagen fiber rearrangement and an increase of collagen fibers deposition and crosslinking [2, 3]. These observations suggest that collagen fiber reorganization favors cancer progression. Understanding the mechanical behavior of fibered structures has also a huge biological interest to produce biomimicking materials within tissue engineering. Reproducing structures with similar properties to the ECM allows *in vitro* studies to resemble *in vivo* material behavior [4, 5] that serves to organize and regulate cells and that may be relevant in developing new therapies for pathologies [6].

Due to the variety of fibered structures and their relevance in many biological processes, there is a widespread interest in the scientific community to understand and characterize the highly non linear mechanical behavior of these networks [7]. Based on high resolution fiber images available in the literature [4, 8–10], fibrils can be found in different curved shapes such as twisted, crimped or non-regular curved fibrils. When fibrils present a curved shape, applying uniaxial deformations will unroll the fibril under a low stress state. Eventually, maintaining the strain rate will lead to a straighter shape of the fibril. As the distance between fibril ends become similar to the fibril length, fibril mechanical behavior shows rigidization [10, 11]. Fibrils suffer higher stress in order to maintain strain rate, showing a linear behavior once fibril is completely straight [12]. This rigidization occurs not only in isolated fibrils. As fibril matrices are stretched, fibrils axial direction is aligned with the stretch direction, leading to the stiffening of the matrix mechanical response [13]. Therefore, this randomly-curved shape in the fiber level (micro level) introduces a non-linear behavior in the stress-strain response at the tissue level (macro level) due to the stiffening associated to fibril unrolling and alignment in the loading direction.

Modeling the mechanical behavior of fibered tissues has been the focus of the development of a wide variety of models (see [14] for a review). The mechanics of the fibril network is dependent of the spatial distribution of the fibers and the constitutive behavior of fibers and crosslinks [14]. An essential input for any fiber network model is the mechanical behavior of a single fiber. Its non-linear fibrous mechanical response can be modeled through either phenomenological or structural models. Phenomenological models are *ad hoc* models that fit fibril rigidization stress-strain curves considering exponential, polynomial or logarithmic functions [15, 16]. In this modeling strategy, a strain energy density function is deployed to express the mechanical property of individual fibers and correlations with experimental results are needed to fit the model parameters [17–21]. Although phenomenological models predict the behavior of the fibers accurately, they do not account for any structural information. On the other hand, fibrils with different shape on their twisted state show different mechanical response in the early stretching phase before the completely straight fibril shape with the same physiological paramaters. As fibril can present randomly-curved shapes, it is interesting to evaluate their structural evolution during the stretching process. Hence, it is possible to use structural models in order to describe explicitly fibril mechanical response, where physiological parameters will give the accurate stress-strain results for a specific fibril shape. This way, differences in fibril mechanical response due only to fibril shape differences are reproducible. Several structural models have been developed for fibrous structures by incorporating the network geometry in the model, either through image processing [22–28] or numerically generated [13, 29–38]. Voronoi tessellations, Delaunay triangulations and other networks are used to generate the random fiber systems and use straight fibrils to replicate the fiber microstructure. Therefore, these models are limited as their non linear constitutive behavior comes from the gradual alignment of the fibers in the direction of the load applied but not from the physiological unrolling of the fibers. Others include waviness with collagen fibers in a load free state through helical springs [39] and predefined curved fibers [40, 41].

In this work, a multiscale formulation approach is used to mechanically characterize the non linear behavior of randomly curved fibered structures from the microstructure. Contrary to previous existing structural models, a fibered matrix model is built from the analysis of the mechanics of a randomly curved single fiber. The isotropic / anisotropic fiber distribution, their evolution during straining and the fluctuations of density are modeled at the micro level. Upon homogenizing these fibered matrices, the macroscopic behavior is captured at the macroscale. Therefore, the key aspect of this study is to recognize the importance of the curved shape of fibered structures on their three dimensional mechanical behavior under tensile, compression and shear loads.

## 2. Single fiber

This section describes the mechanical analysis of a single fiber, from the generation of randomly curved single fibers, to the mathematical analysis of their mechanical behavior at tensile and compression tests.

### 2.1. Fiber generator

Here we present the methodology to build different 3D randomly curved collagen fibers (see Fig. 1). We define the input parameter *L* as the distance between both ends of the fiber. This distance is discretized into *n* points, in which an auxiliary polar system (*r, θ*) is defined. The polar angle is randomly selected from a uniform distribution such that *θ* ∈ [0, 2*π*). Analogously, the fiber eccentricity from the main axis, *r* is selected from a uniform distribution *r* ∈ [0, 2*R*] (see Fig. 1a). Therefore, the coordinates of the different discretized points along the fiber are given as:

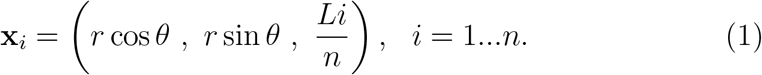

The parameter *n* represents in Eq. (1), the number of twists of the curved fiber along its length. On the other hand, *R* is defined from Fig. 1a as follows:

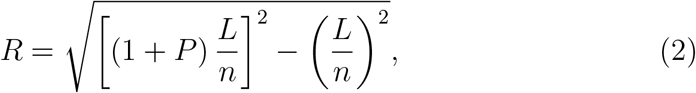

with *P* (*P* ≥ 0) being a relative length parameter of the fiber length versus the initial length of the end points.

**Figure 1.**
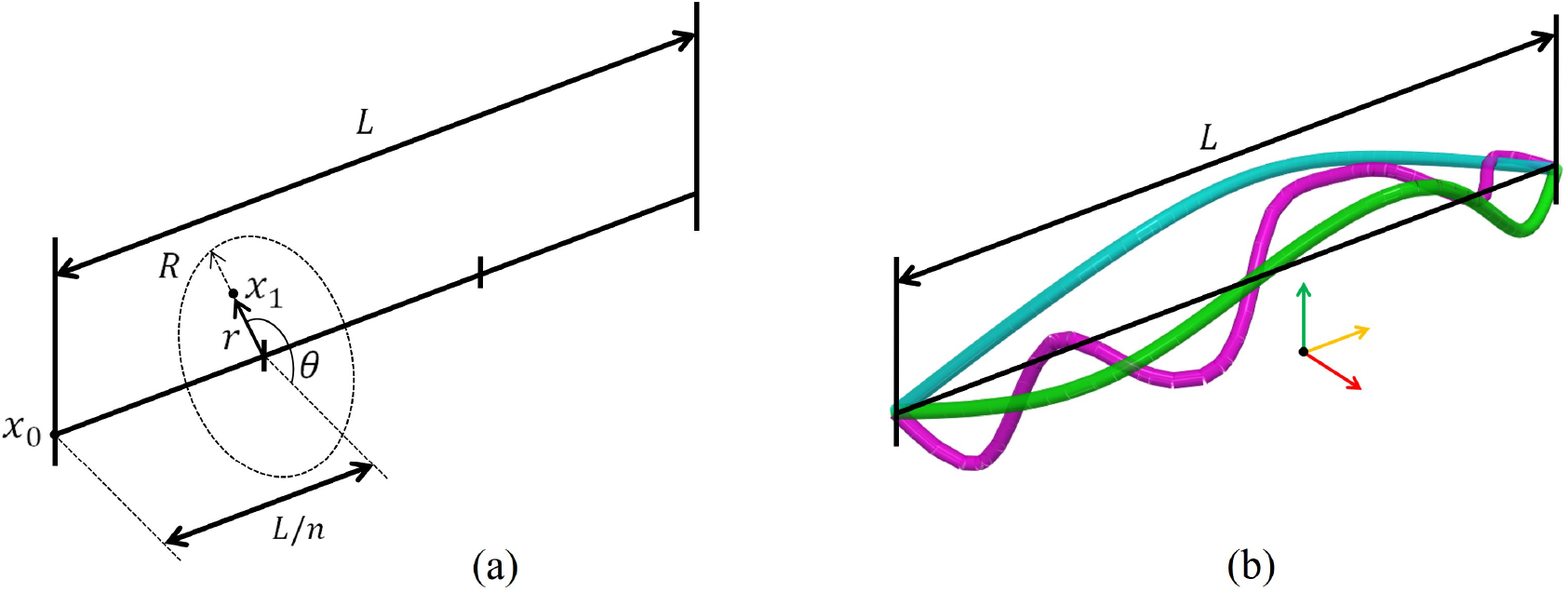
Schematics of the fiber generator. (a) Definition of the discrete points ***x***_*i*_ (for *n* = 3). (b) Different individual randomly-curved fiber in 3D representation, for input parameters *n* = 3, 5 and 10; and *L*_*f*_ */L* = 1.11, 1.21 and 1.4 for blue, green and magenta fibers, respectively.

Once the 3-D spatial points ***x***_*i*_ have been created (coarse discretization of the fiber), the curved fiber is smoothed by defining a spline through points ***x***_*i*_. This curve is then discretized (fine discretization of the fiber) into *NE* elements. The final length of the fiber *L*_*f*_ is then defined as 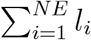, being *l*_*i*_ the element length. Different generated individual fibers can be seen in Fig. 1b.

### 2.2. Mathematical formulation

The generated curved fiber is discretized into *NE* Euler-Bernouilli beam finite elements. Each element contains 2 nodes and 6 degrees of freedom (3 translations and 3 rotations) per node. We follow an updated lagrangian formulation such that the fiber geometry is updated at each current time *t* of analysis:

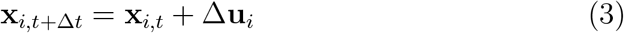

with **x**_*i,t*+Δ*t*_ and **x**_*i,t*_ the position vectors of node *i* at configurations *t* + Δ*t* and *t*, respectively, and Δ**u**_*i*_ the vector of displacements of node *i* from configuration *t* to *t* +Δ*t*. This vector Δ**u**_*i*_ is obtained after the discretization and assembly of global vector and matrix following a structural matrix finite element analysis [42]:

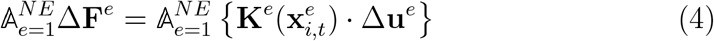

being 𝔸 the assembly operator; and Δ**F**^*e*^ and Δ**u**^*e*^ the incremental nodal vector of structural forces and displacements/rotations at element *e*, respectively. 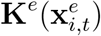 is the matrix of element *e* computed in the configuration *t* with element nodal coordinates 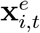. After assembly, the (global) system in Eq. (4) yields,

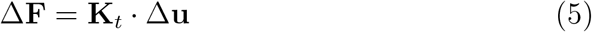

The total nodal force vector is then updated as:

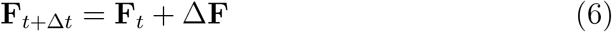

The finite element implementation of Eq. (5) was developed in Matlab@. The computer implementation was validated in Fig. 2 versus the exact solution of a nonlinear cantilever rod subjected to a point bending moment.

**Figure 2.**
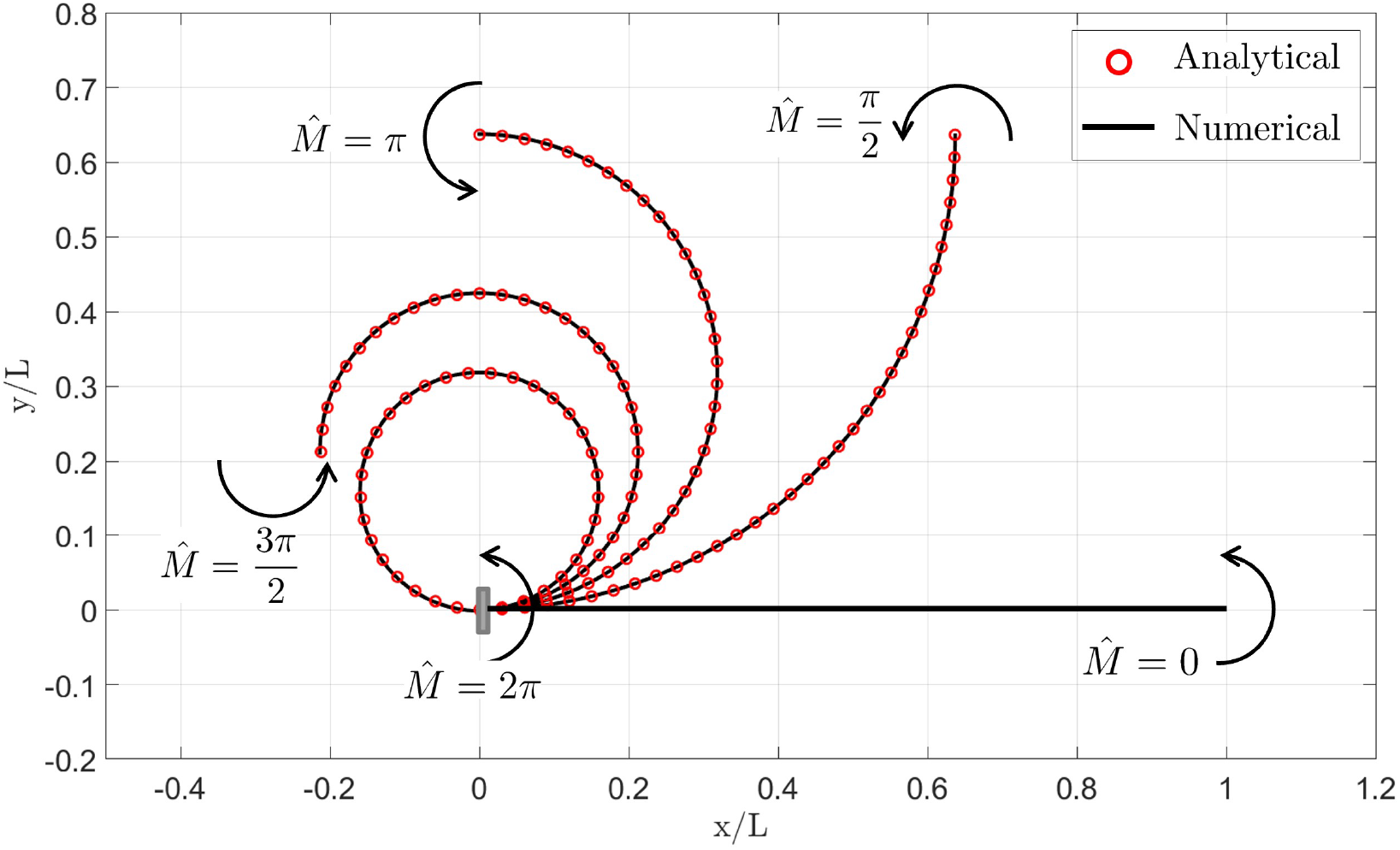
Exact solution of a nonlinear cantilever rod subjected to a point dimensionless bending moment 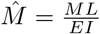 versus its finite element implementation. The analytical solution is an arc with dimensionless radius 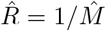 [43]. The rod was discretized into 150 elements, and the moment step was 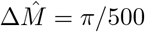.

### 2.3. Results

In this section, we analyze the tensile/compression behavior of a single randomly curved fiber for a range of model parameters, namely, fiber end points length *L*, fiber diameter *d*, fiber elastic modulus *E*; and fiber length to fiber end points length ratio *L*_*f*_ */L* (being this parameter an indication of the wavy/curly shape of the fiber). A baseline case is selected, with parameters *L* = 20 *μm, d* = 0.2 *μm, E* = 100 MPa and *L*_*f*_ */L* = 1.1. These parameters are taken as an estimation of the order of magnitude found in the literature for collagen (see Table 1).

**Table 1:** Range of parameters of natural collagen structures available in the literature. Dispersion of parameters might be due to different animal origin.

| Property | Value | References |
| --- | --- | --- |
| Linear modulus (MPa) | 110-1470 | [44] |
|  | 0-32 | [45] |
|  | 2000-7000 | [12] |
| Fiber length ($\mu\text{m}$) | 100-200 | [12] |
|  | 6-18 | [44] |
|  | 10-100 | [10] |
| Fiber diameter ($\mu\text{m}$) | 0.1-0.5 | [12] |
|  | 0.418-0.446 | [4] |
|  | 0.01-0.25 | [46] |
|  | 0.205-0.905 | [44] |

Figure 3 shows the mechanical behavior of a single fiber in uniaxial stretch-stress tests for the range of analyzed model parameters. The tensile region (*λ >* 1) of the curves in Fig. 3 shows a nonlinear behavior divided [43]. The rod was discretized into 150 in two different regimes: first a nonlinear part followed by a strain stiffening due to fiber straightening. This behavior can be seen in video S1 of the supplementary material, for the baseline fiber. Interestingly, it can be seen in the compression region (*λ <* 1) in Fig. 3, that our formulation naturally recovers the instability (buckling) phenomenon developed by the fiber at compression.

**Figure 3.**
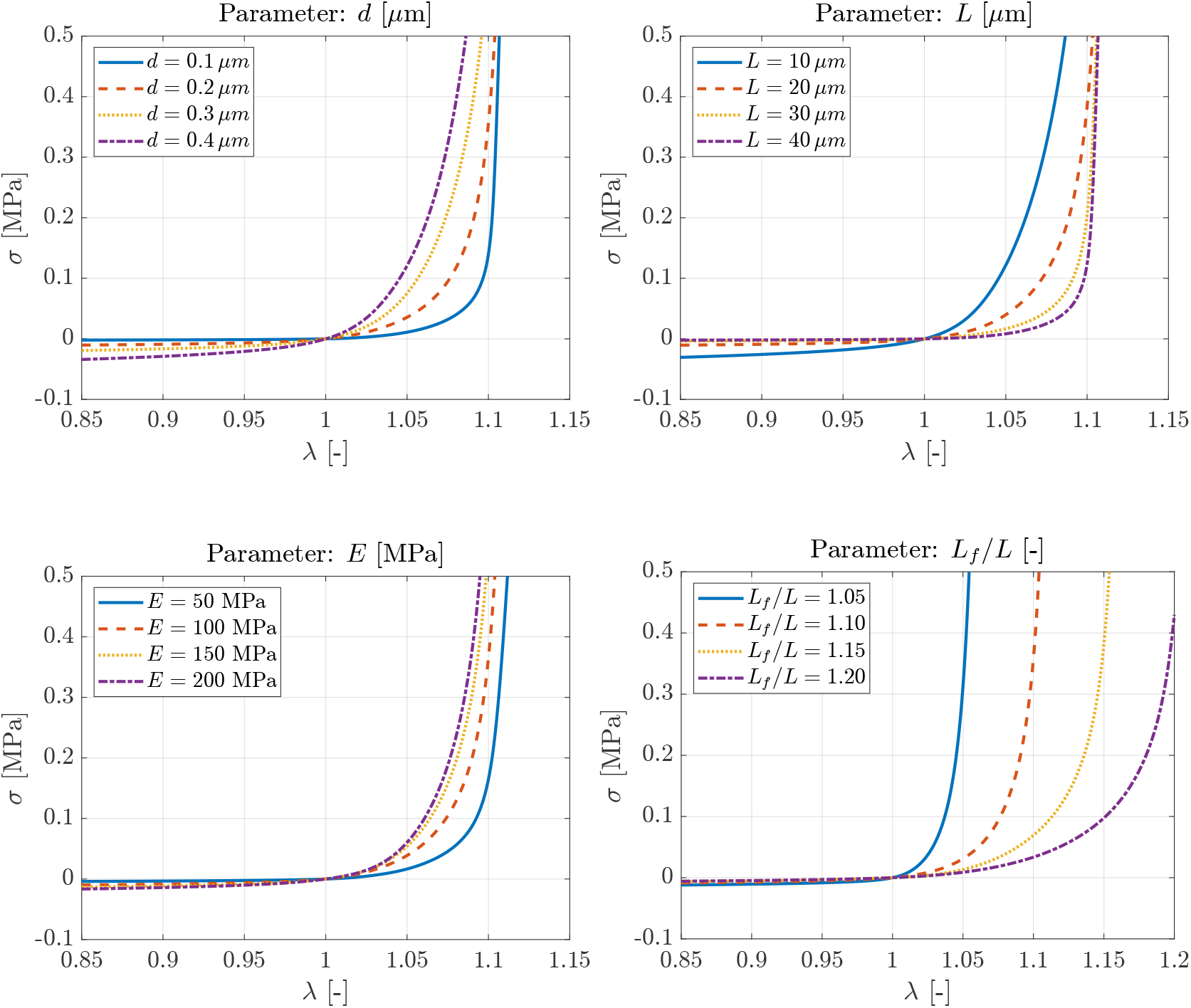
Parametric analysis of the mechanical behavior of elastic single randomly curved fiber. Model parameters are varied within the range found in the literature (see Table 1), from a baseline case that with parameters *L* = 20 *μm, d* = 0.2 *μm, E* = 100 MPa and *L*_*f*_ */L* = 1.1.

On the other hand, the incremental elastic energy of the fiber can be defined as follows:

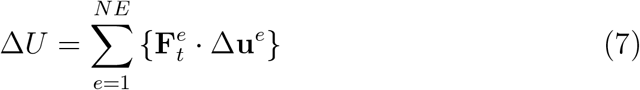

The total accumulated elastic energy of the fiber is then obtained as:

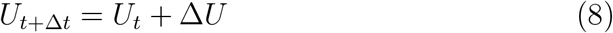

The total accumulated elastic energy of the fiber is split from Eq. (8) into axial, bending an torsional energies as follows:

- Axial energy is obtained by selecting the axial forces and axial displacements product components in Eq. (8).
- Bending energy is obtained by selecting the shear forces/bending moments and shear displacements/bending rotations product components in Eq. (8).
- Torsional energy is obtained by selecting the torque moment and torque rotation product components in Eq. (8).

Figure 4 shows the different fiber accumulated energies for the uniaxial tensile/compression stretch-stress tests of the baseline single fiber. Figure 5 additionally plots the different fiber accumulated energies relative to the total energy. According to Fig. 5, the first tensile region of the curve (represented by point 1) is governed mostly by fiber bending (with minor contribution of the axial and torsional energies), up to a certain stretch (point 2) in which the fiber is almost straight except a small portion of the fiber. Indeed, the deformation point 2 represents the onset of straightening of the fiber, according to the overall length of the fiber given by parameter *L*_*f*_ */L* (equal to 1.1 in this case). Once the fiber is completely straight (point 3) the most important contribution of energy is due to axial deformation. In the compression regime (points 4 and 5) the most important contribution is the bending energy as the fiber is getting wavy (curly). The torsional energy is almost negligible in the different regimes of deformation of the analyzed fiber. It was checked (data not shown) that the total accumulated elastic internal energy of the fiber, obtained from Eqs. (7)-(8), is equal to the external energy developed by the applied force in the fiber. Note that the point *λ* = 1 was not computed in Figure 5, that all the energies are null, to avoid this singularity.

**Figure 4.**
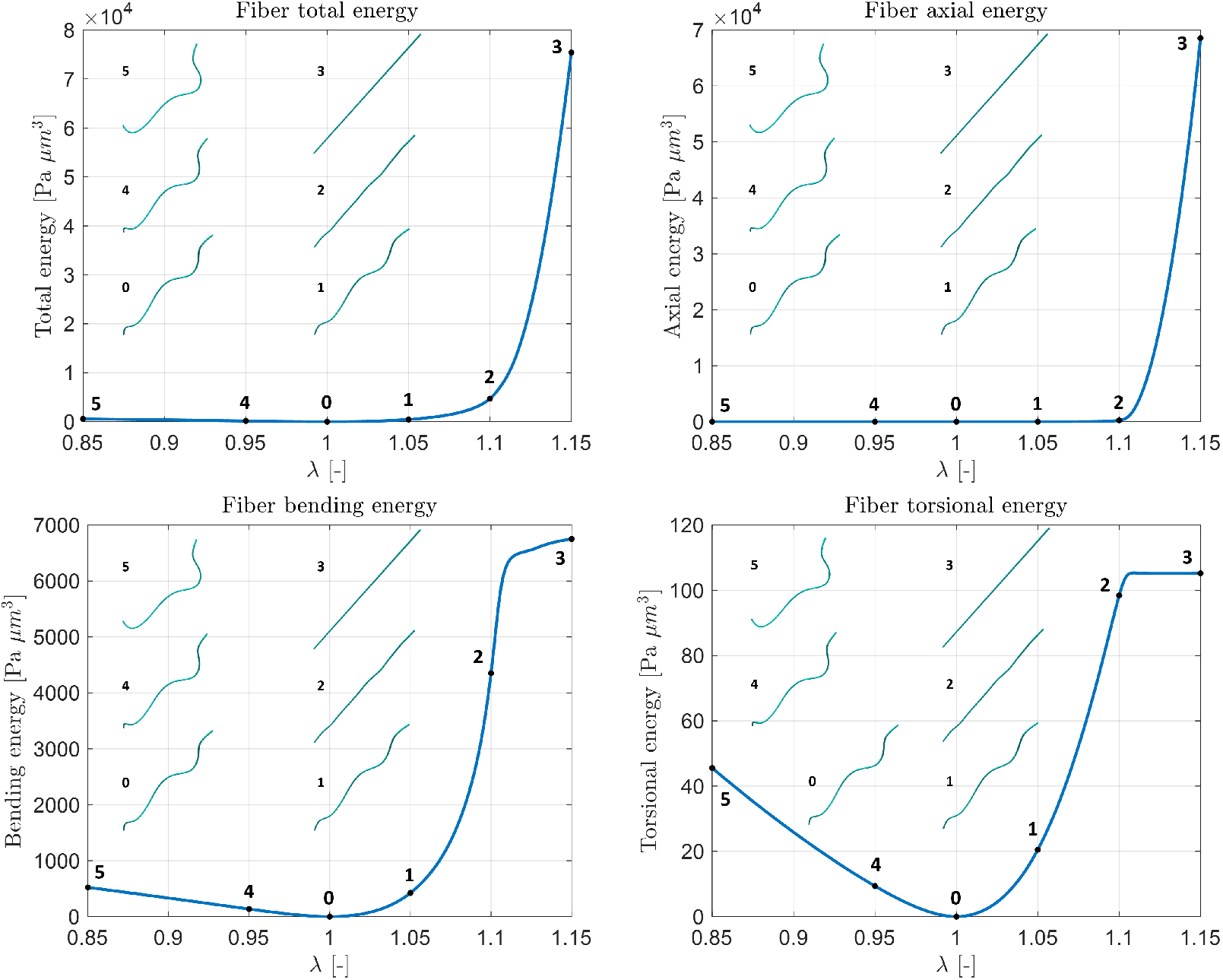
Total accumulated energy, as well as axial, bending and torsional components of the accumulated energy for a single randomly curved fiber with parameters *L* = 20 *μm, d* = 0.2 *μm, E* = 100 MPa and *L*_*f*_ */L* = 1.1. Inset fibers in the figures represent the deformation of the fiber for the selected stretch points.

**Figure 5.**
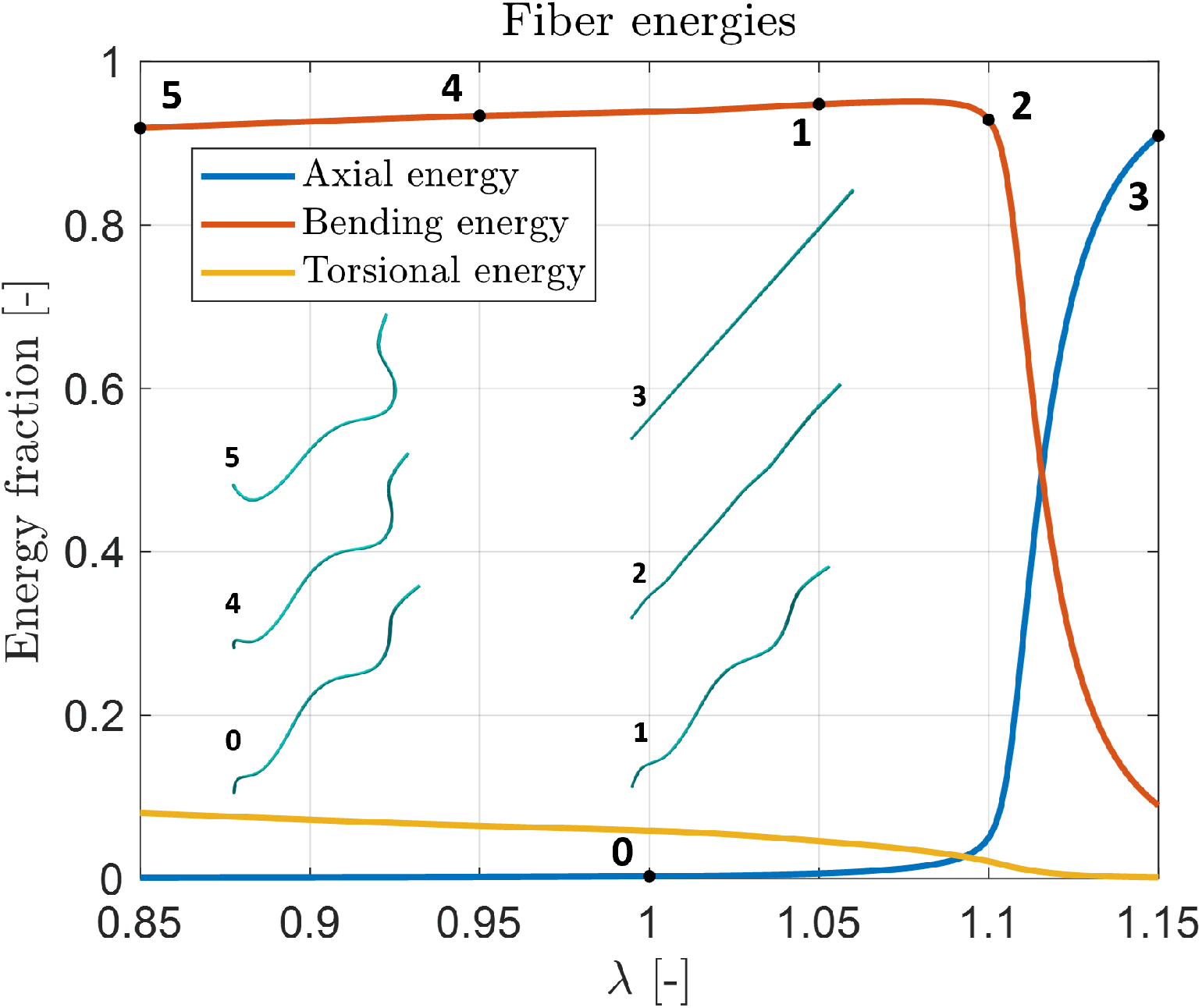
Axial, bending and torsional accumulated energies relative to the total energy, for a single randomly curved fiber with parameters *L* = 20 *μm, d* = 0.2 *μm, E* = 100 MPa and *L*_*f*_ */L* = 1.1. Inset fibers in the figures represent the deformation of the fiber for the selected stretch points.

Keeping in mind that the first tensile/compression regions of the stretch-stress curve of the fiber behavior is governed by bending, and the strain stiffening region by the axial energy, the impact of model parameters shown in Fig. 3 can be easily understood. Consistently, we can see a stiffer behavior (both at tensile and compression) for increasing fiber stiffness (*E*) and diameter (*d*); and decreasing length (*L*). Indeed, in qualitative terms, the fiber behavior depends as ∝ *Ed*^4^*/L*^3^ in the bending region; and as ∝ *Ed*^2^*/L* in the axial (strain stiffening) region. The effect of fiber wavy/curly (parameter *L*_*f*_ */L*) is seen in Fig. 3 as a stiffer behavior for decreasing *L*_*f*_ */L* (less wavy fiber) with shorter bending region and, consequently, with faster development of the strain stiffening region. Indeed, the onset of the development of the strain stiffening region is correlated with parameter *L*_*f*_ */L*.

## 3. Fibered matrix

This section describes the multiscale mechanical analysis of a fibered matrix, composed of randomly curved fibers, from the mechanical interaction of fibers in a representative volume element (RVE).

### 3.1. Matrix generator

The generation of the RVE is established under the basis of the definition of RVE, that is a selected volume of the fiber microstructure that statistically represents the heterogeneity of the matrix [47]. Our RVE is composed of isolated fibers *Nf*_*i*_ and crosslink fibers *Nf*_*x*_, such that the total number of fibers of the RVE is *Nf* = *Nf*_*i*_ + *Nf*_*x*_. Therefore, the volume concentration of fibers is defined as *V*_*c*_ = *Nf/V*_*RVE*_, with *V*_*RVE*_ being the volume of the RVE. The algorithm to generate fibered matrices proceed as follows:

#### Box 1

Algorithm to generate fibered matrices

0. Set the volume concentration of fibers *V*_*c*_ and the volume of the RVE, and consequently, *Nf, Nf*_*i*_ and *Nf*_*x*_.

1. Generate *Nf*_*i*_ isolated fibers (see section 2.1).
2. FOR *m = 1*..*Nf*_*i*_
  2.1 Randomly select *N*_*mx*_ crosslinking nodes of fiber *m*.
  2.2 FOR *j* = 1..*N*_*mx*_

i. Create a set with potential connecting candidate crosslinking nodes *k*, for fibers *n* = 1..*Nf*_*i*_ (*n* ≠ *m*) such that *L*(1 − *ε*) ≤ *L*_*j−k*_ ≤*L*(1 + *ε*).
ii. Randomly select node *j* from the set.
iii. Generate a crosslinking fiber from nodes *j*−*k* (with length *L*) as in section 2.1. END FOR. END FOR.

We set *N*_*mx*_ = 1 in Box 1, meaning that a crosslinking fiber is generated per isolated fiber. Therefore, *N*_*mx*_ represents the degree of crosslinking of the matrix. On the other hand, the endpoints length of the crosslinking fiber is set to the same length *L* of the isolated fiber, according to item *i* in Box 1, up to a tolerance *E* which is set to 0.01 in our code.

### 3.2. Multiscale formulation

The overall macroscopic mechanical behavior of a fibered matrix and its evolution, is obtained from the micromechanical interaction of fibers within the RVE in a multiscale fashion (see Fig. 6). In the macroscale, we define the (Cauchy) stress tensor and the (logarithmic) strain tensor as 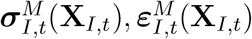, respectively, in a macroscopic (Gauss) material point *I* of the macroscale (with coordinates **X**_*I,t*_), for load time *t* (see Fig. 6). On the other hand, in the microscale, we define the variables 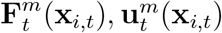 associated to an Euler-Bernoulli beam in a material point (node) *i* of the microscale (with coordinates **x**_*i,t*_) for load time *t*. 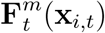 represents the vector of nodal structural forces (forces and moments), whilst 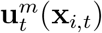 is the vector that contains nodal displacements and rotations (see Fig. 6).

**Figure 6.**
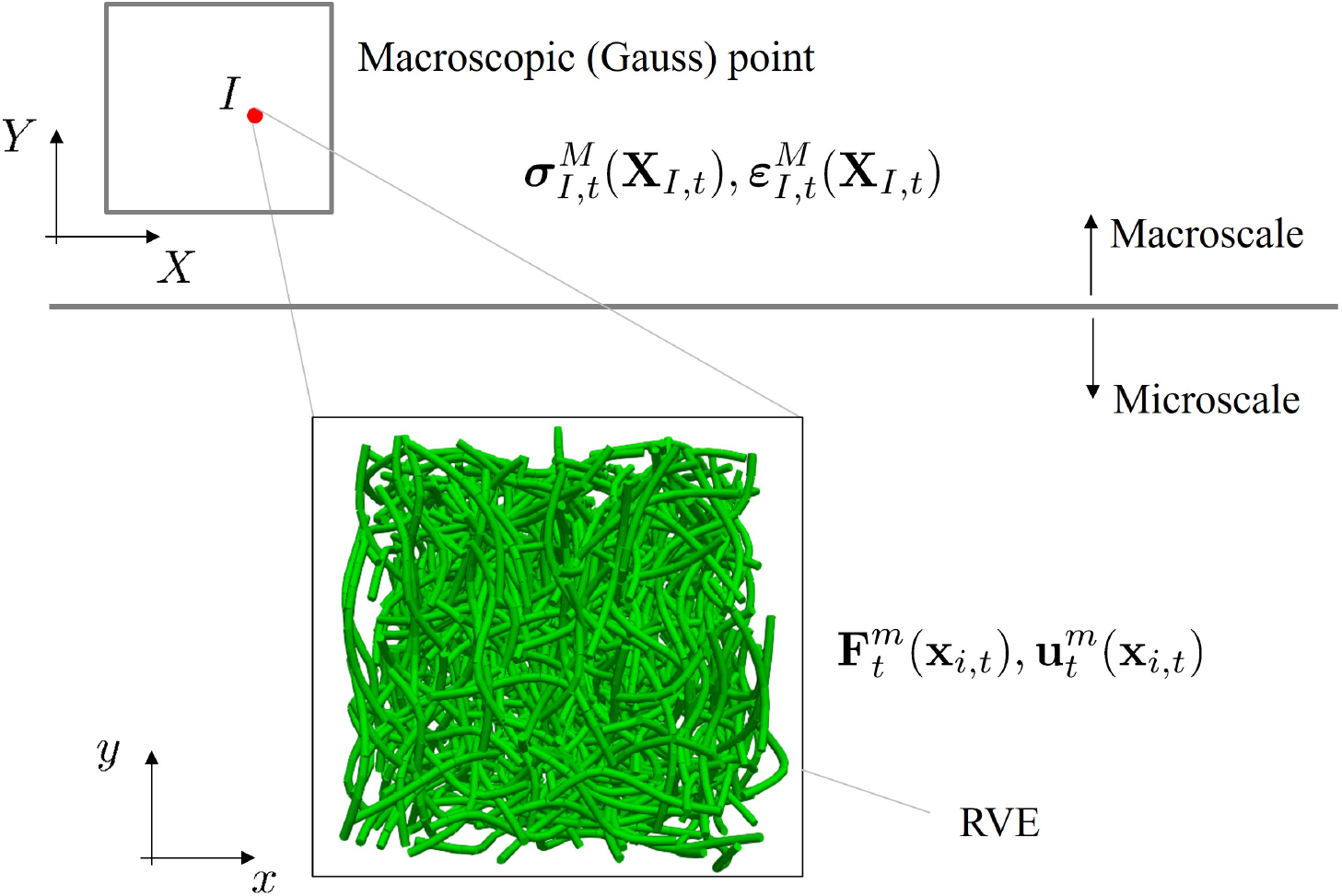
Schematics of the proposed multiscale formulation of fibered matrices.

As stated in the analysis of the single fiber, we follow an updated lagrangian approach such that the configuration (finite element mesh) is updated at each load time *t*. Consequently, at the macroscale, the logarithmic stain tensor is obtained by integrating the strain rate, numerically, in a material frame of reference:

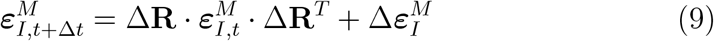

where 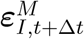 and 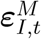 are the total strains at increments *t* +Δ*t* and *t*, respectively; Δ**R** is the incremental rotation tensor; and 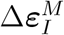 is the total strain increment from increment *t* to *t* + Δ*t*. Furthermore, the macroscopic strain density energy from increment *t* to *t* + Δ*t* is defined as:

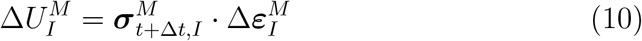

Analogously, the kinematics at the microscale is defined as:

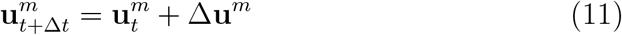

where 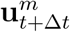 and 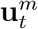 are the total displacements/rotations vectors at increments *t*+Δ*t* and *t*, respectively; and Δ**u**^*m*^ is the total displacement increment vector from increment *t* to *t* + Δ*t*. The displacements increment vector is defined at nodes *k* of the microscale through the strain increment at the macroscale as:

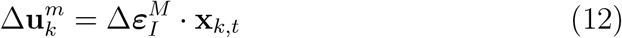

where ***x***_*k,t*_ are the (updated) microscopic coordinates, at load time *t*, of the points *k* in which the macroscopic strain is affinely transmitted to the microscale. We assume that points *k* are the endpoints (knots) of the fibers. On the other hand, the global nodal forces vector is updated as follows,

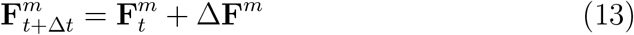

Using Eq. (5) in (13) yields:

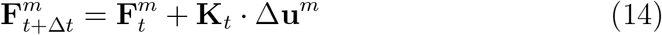

Finally, the total microscopic strain density energy from increment *t* to *t* + Δ*t* is defined as:

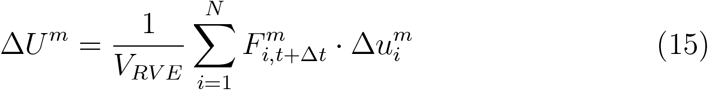

where *N* is the number of nodes of the microstructure. On the other hand, we use the Hill’s equality of energies from the transition from the macro to the microscale [47, 48]

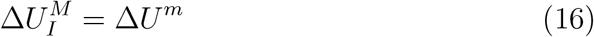

Thus, using (10) and (15) in (16) yields,

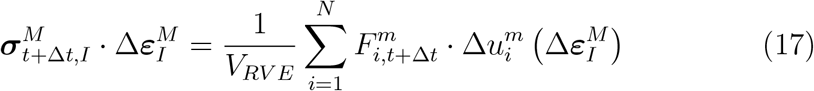

Note that in Eq. (15), 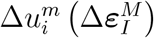 represents the incremental nodal displacements/rotations of the microstructure, as a result of the solution of the discrete finite element system Δ**F**^*m*^ = **K**_*t*_·Δ**u**^*m*^ with prescribed displacements at nodes *k* according to Eq. (12). Therefore, the different *kl* components of the stress tensor 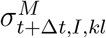 are obtained using Eq. (17) as follows:

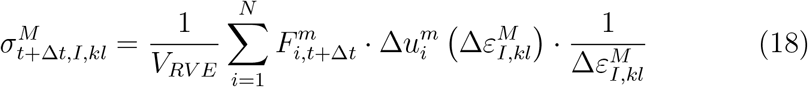

Eq. (18) is equivalent to,

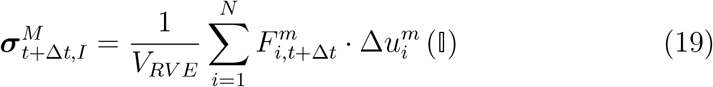

with 𝕀 being the unit tensor:

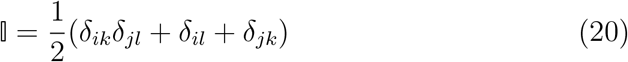

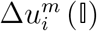 represents the incremental nodal displacements/rotations of the microstructure, as a result of the solution of the discrete finite element system Δ**F**^*m*^ = **K**_*t*_ · Δ**u**^*m*^ with prescribed displacements at nodes *k* for unit strain tensors, following Eq. (12).

Using the decomposition of the nodal forces vector (13) in (18), the incremental stress tensor can be obtained as:

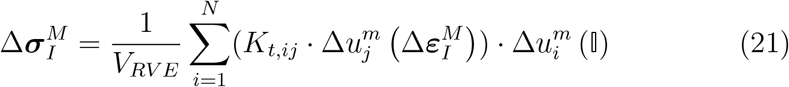

The macroscopic tangent stiffness tensor can be then obtained as:

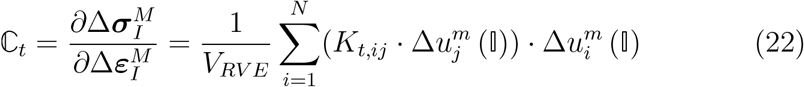

Box 2 summarizes the main steps of the developed multiscale algorithm.

#### Box 2

Algorithm to compute macroscopic stress and strain quantities from microscopic fibered matrices.

1. **Macroscale**: 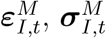available at load increment *t*.
2. **Macroscale:** Set next strain increment 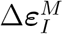.
3. **Microscale:** Obtain 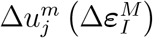 and 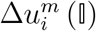 from result of the solution of the system Δ**F**^*m*^ = **K**_*t*_ · Δ**u**^*m*^ with prescribed displacements at nodes *k* for strain tensor 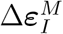 and unit strain tensors 𝕀, respectively, following Eq. (12).
4. **Microscale:** Update microscopic coordinates as 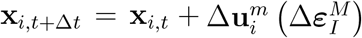.
5. **Macroscale:** Compute 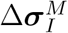 according to Eq. (21). Compute the macroscopic tangent stiffness tensor ℂ according to Eq. (22).
6. **Macroscale:** Update macroscopic quantities as 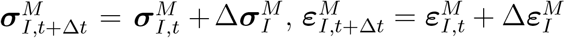
7. *t* ← *t* + Δ*t*. GO TO 1.

Finally, the incremental elastic internal microscopic energy of a fiber *f* of the microstructure of the matrix, Δ*U*^*m,f*^, is computed as,

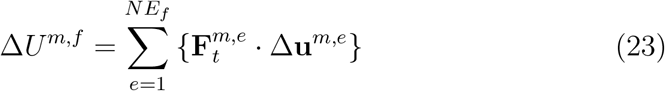

where 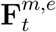 and Δ**u**^*m,e*^ are the total structural forces and displacements/rotations nodal vectors at element *e*, respectively. *NE*_*f*_ is the number of elements of the fiber *f*. The total accumulated elastic energy of each fiber *f* of the microstructure of the matrix is obtained as:

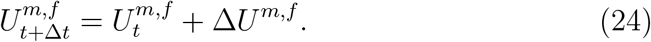

On the other hand, the total accumulated microscopic strain density energy is computed as,

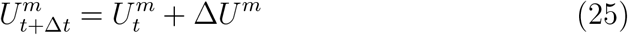

with Δ*U*^*m*^ obtained from Eq. (15).

### 3.3. Results: Tensile/Compression

In this section, different fibered microstructures—with different microsctructural characteristics—are subjected to a macroscopic tensile and compression stress state. The stress-stretch curves of the virtual tests are then obtained following the multiscale approach exposed in section 3.2.

For this purpose, a vertical stretch velocity 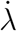 (*Y* -direction) is prescribed. The macroscopic incremental strain tensor (point 2 of Box 2 of the multiscale formulation) is obtained from the solution of:

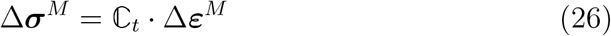

Δ***ε***^*M*^ is obtained from the solution of the system in Eq. (26) with uniaxial (mixed boundary conditions) stress state. In this sense, 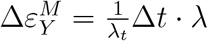 for the macroscopic incremental strain tensor, and 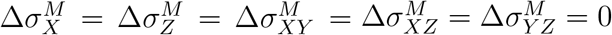.

Figure 7 shows the mechanical behavior of the considered matrices in uniaxial stretch-stress tests. In this figure, different randomly curved fibered matrices, at different fiber versus volume concentration (parameter *V*_*c*_), were generated for the range of analyzed model parameters. These parameters were varied within the order of magnitude of real biological fibered matrices (see Table 1), from a baseline case with parameters *L* = 30 *μm, d* = 0.4 *μm, E* = 150 MPa, *L*_*f*_ */L* = 1.15 and *V*_*c*_ = 2.05 · 10^6^ fibers*/mm*^3^.

**Figure 7.**
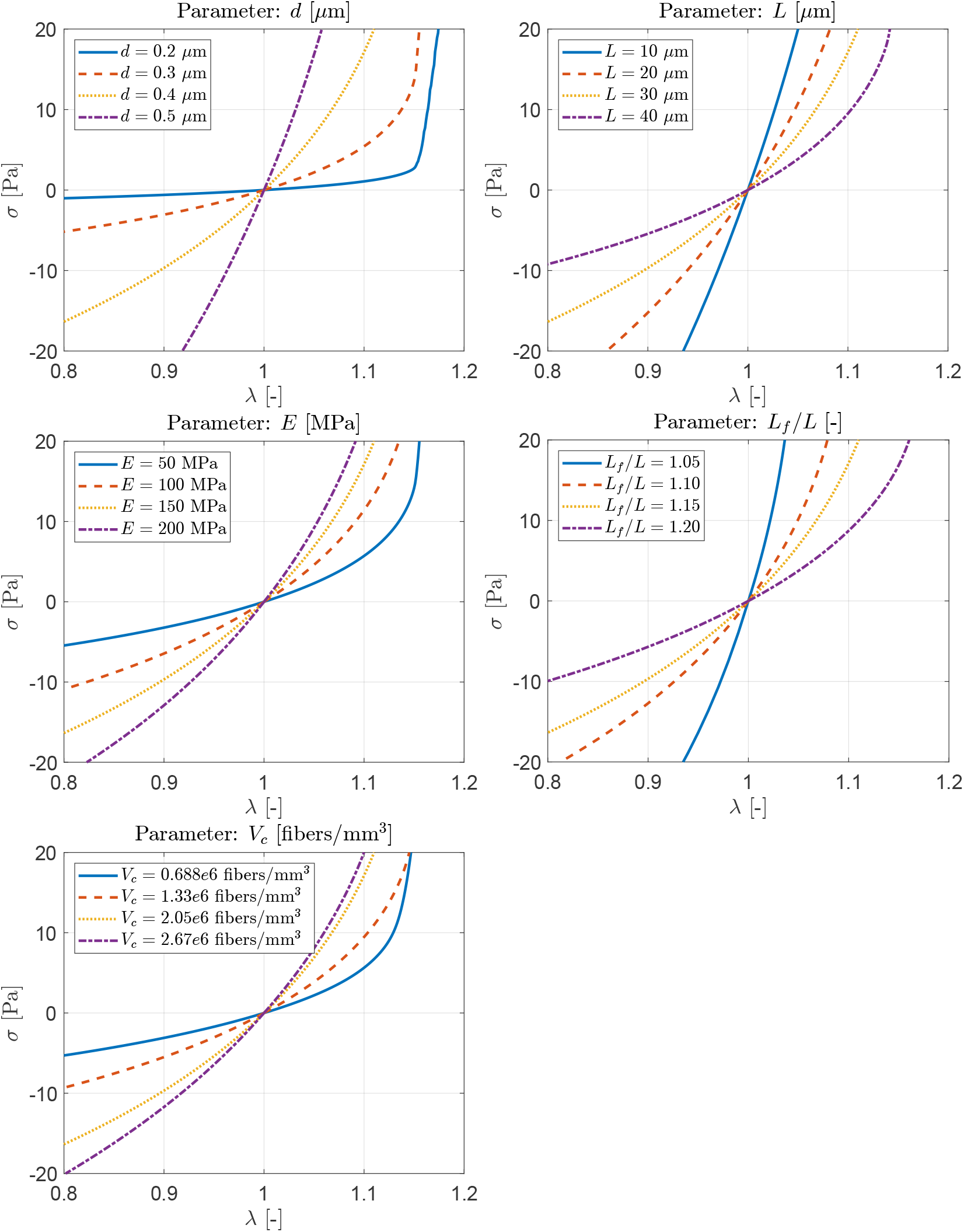
Tensile/compression tests. Parametric analysis of the mechanical behavior of elastic randomly curved fibered matrices. Model parameters are varied within the range found in the literature (see Table 1), from a baseline case with parameters *L* = 30 *μm, d* = 0.4 *μm, E* = 150 MPa, *L*_*f*_ */L* = 1.15 an4d1*V*_*c*_ = 2.05 · 10^6^ fibers*/mm*^3^.

Similarly to the single fiber, the tensile region (*λ >* 1) of the curves in Fig. 7 shows a linear behavior at small strains followed by a nonlinear behavior. In this case, the nonlinear regime is the consequence of both fiber straightening and fiber alignment with the direction of load. Besides, the loss of stiffness due to the instability (buckling) phenomenon developed by the fibers at compression, can be seen in the compression region (*λ <* 1) in Fig. 7. According to Figure 7, and analogously to the single fiber behavior, the overall tensile/compression behavior of the matrices is stiffer for increasing fiber stiffness (*E*), fiber diameter (*d*) and concentration of fibers. On the other hand, the overall tensile/compression behavior of the matrices is softer for increasing fiber length (*L*) and fiber curvature (*L/L*_*f*_).

The total accumulated strain density energy developed by the microstructure, computed from Eq. (25), can be seen in Figure 8 for the uniaxial tensile/compression stretch-stress test. Results are therefore shown as the (accumulated) microscopic contribution of all the fibers of the matrix along the deformation path, for the considered baseline matrix defined above. Note that this quantity is equal to the total accumulated strain density energy developed at the macrostructure (Eq. (10)), by virtue of Eq. (16). In Figure 8, the axial, bending and torsional components of the total accumulated strain density energy are also shown. These quantities were obtained from Eq. (25), after selecting the axial, bending and torsional components similarly to the single fiber case (section 2.3). As in the single fiber case, it can be seen in this figure the asymmetry of the total strain density energy at tensile (*λ >* 1) versus the compression regime (*λ <* 1). Indeed, matrix compression induce fiber instabilities (buckling) whereas matrix tensile produces fiber unrolling and fiber alignment with load (strain stiffening). Figure 8 additionally plots the different accumulated strain density energies relative to the total strain energy energy.

**Figure 8.**
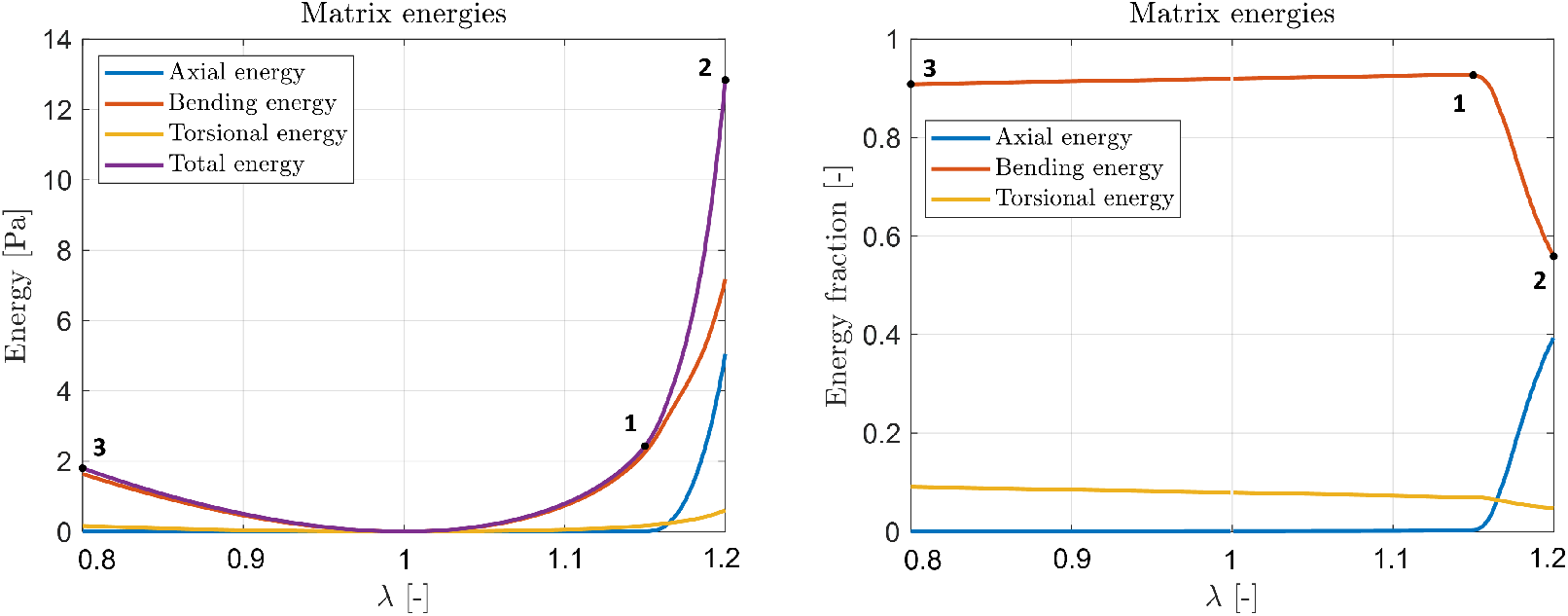
Multiscale energies: Macroscopic. Total, axial, bending and torsional accumulated strain density energies developed by the microstructure for tensile/compression tests of the baseline matrix.

On the other hand, Figure 9 shows the accumulated multiscale energies in the fibers of the microstructure of the matrix for different tensile/compression deformation points, as defined in Figure 8.These quantities were computed from Eq. (24) after selecting the axial, bending and torsional components. The tensile/compression behavior of the baseline matrix, the evolution of its microstructure, and both the axial, bending and torsional accumulated energies in the fibers; can be seen in videos S2, S3 and S4 of the supplementary material. It can be seen in Figures 8 that bending energy dominates in all the analyzed deformation regime, except the strain stiffening region. At the tensile region (*λ >* 1) this energy in employed to fiber unrolling and fiber alignment with load, as corroborated in the microstructure evolution by Figure 9 and video S3. At the compression region (*λ <* 1) the bending energy is employed to fiber buckling, as corroborated in the microstructure evolution (see also Figure 9 and video S3). Moreover, the torsional energy is almost negligible in all the analyzed deformation range, according to Figures 8 and 9.

**Figure 9.**
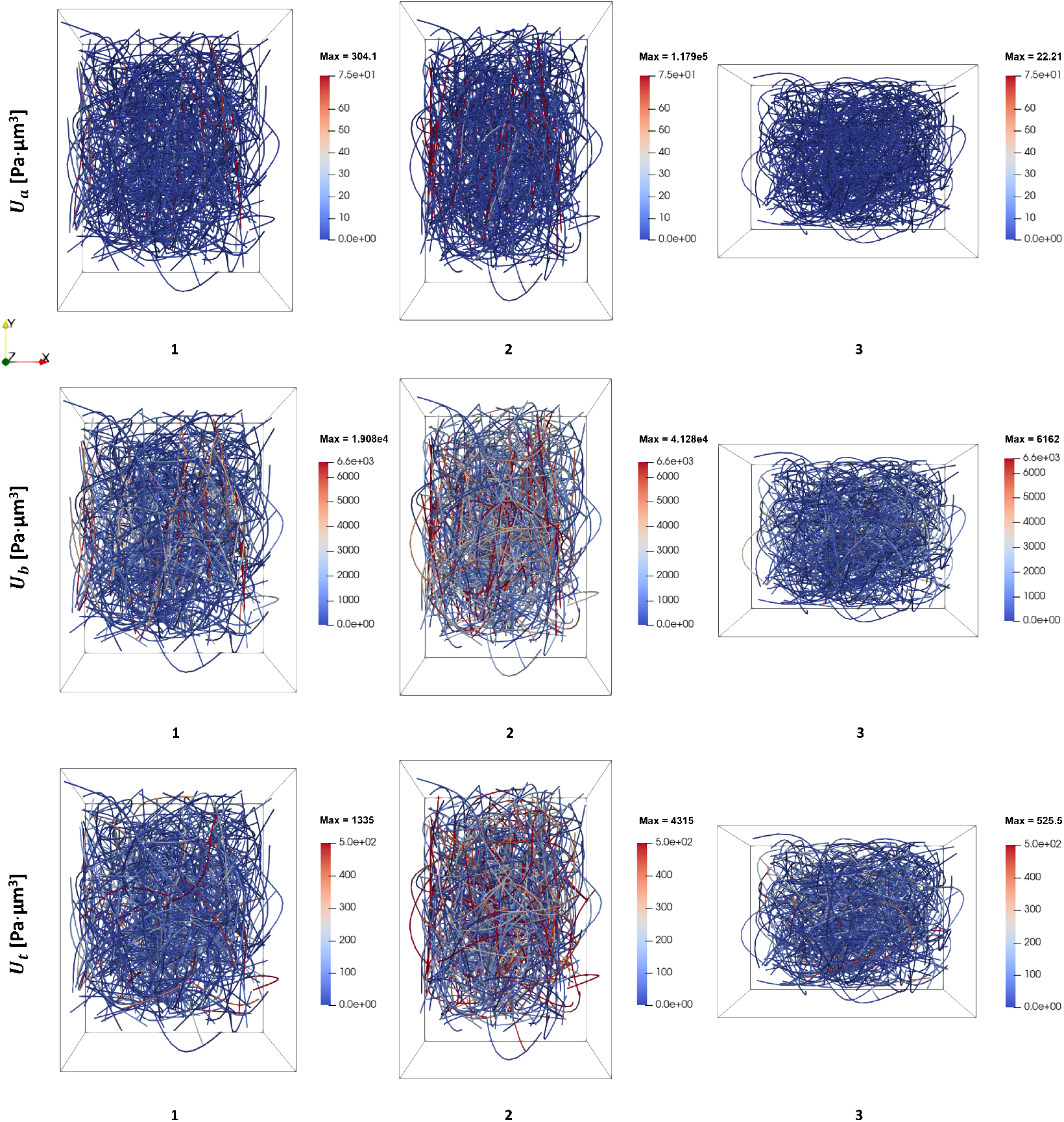
Multiscale energies: Microscopic (fiber scale). Plot in the microstructure of the axial (first row), bending (second row) and torsional (third row) of the accumulated energies in the fibers of the baseline matrix for tensile/compression tests. Results are shown at different deformation points (1,2,3) defined in Figure 8.

### 3.4. Results: Simple shear

Analogously to the previous section, different fibered microstructures are subjected to a macroscopic simple shear stress state. In this case, a shear strain velocity 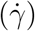 is prescribed, such that the macroscopic incremental strain tensor (point 2 of Box 2 of the multiscale formulation) is fully prescribed as a null value for all components except for 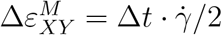.

Figure 10 shows the mechanical behavior of the considered matrices in simple shear stress tests. In this figure, we assumed the same range of variation of the parameters, as well as the same baseline case, than the tensile/compression tests. Therefore, the same microstructures (RVEs) defined for the tensile/compression tests in the previous section, are used here to simulate simple shear tests.

**Figure 10.**
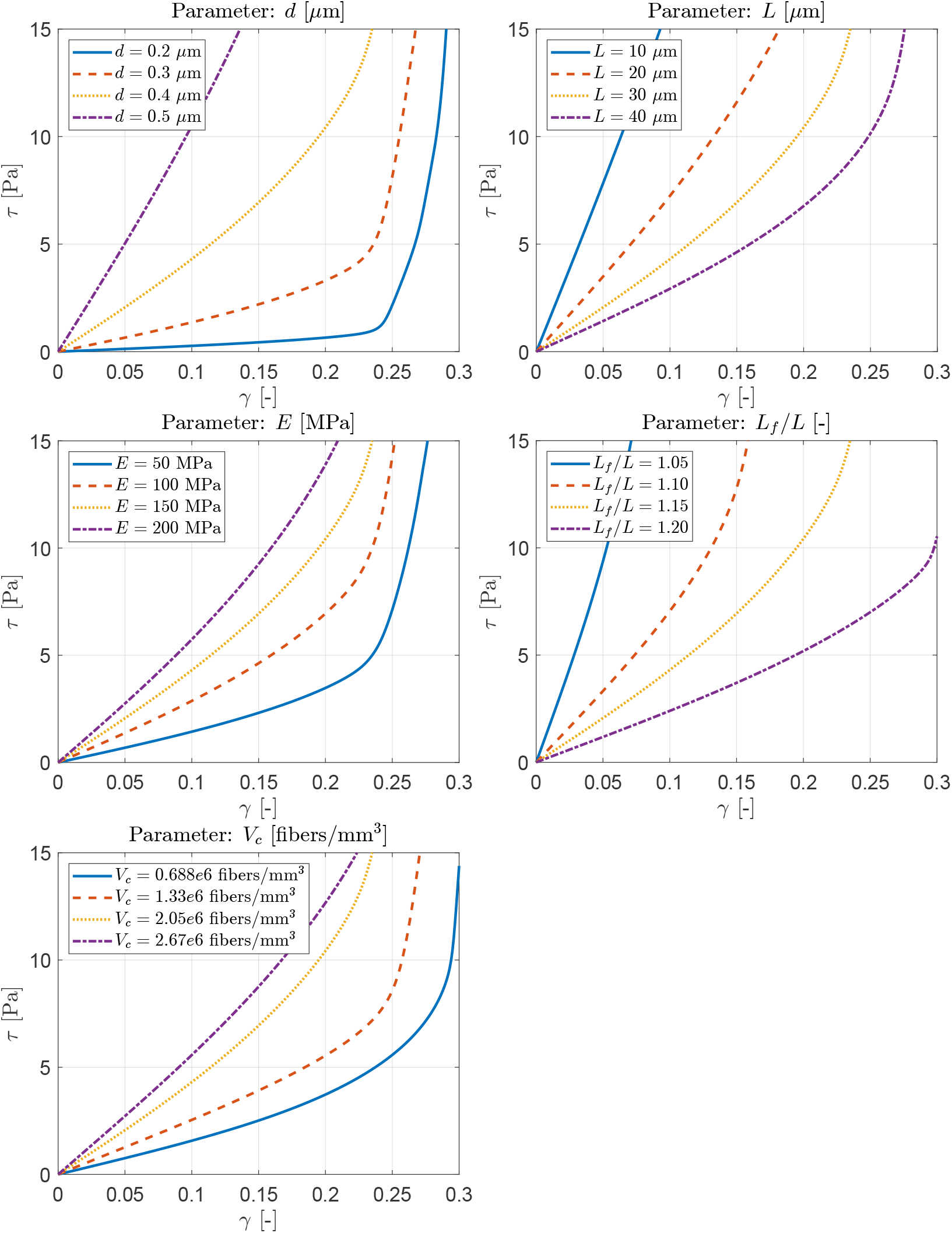
Simple shear tests. Parametric analysis of the mechanical behavior of elastic randomly curved fibered matrices. Model parameters are varied within the range found in the literature (see Table 1), from a baseline case that with parameters *L* = 30 *μm, d* = 0.4 *μm, E* = 150 MPa, *L*_*f*_ */L* = 1.15 and *V*_*c*_ = 2.05 · 10^6^ fibers*/mm*^3^.

The shear behavior of the (virtually) tested matrices, Figure 10, shows also a nonlinear profile. First, a linear region at small strains followed by a nonlinear region, also as a consequence of both fiber straightening (unrolling) and fiber alignment with the direction of load. Analogously to the tensile tests, the shear behavior of the matrices is stiffer for increasing fiber stiffness (*E*), fiber diameter (*d*) and concentration of fibers. On the other hand, the overall tensile/compression behavior of the matrices is softer for increasing fiber length (*L*) and fiber curvature (*L/L*_*f*_).

As in the previous section, the accumulated strain density energies developed by the microstructure for the shear test can be seen in Figure 11. In this figure, results are shown as the (accumulated) microscopic contribution of all the fibers of the matrix along the deformation path, for the considered baseline matrix. The axial, bending and torsional components of the strain density energies were computed analogously to the previous section. Figure 11 additionally plots the different accumulated strain density energies relative to the total strain energy energy.

**Figure 11.**
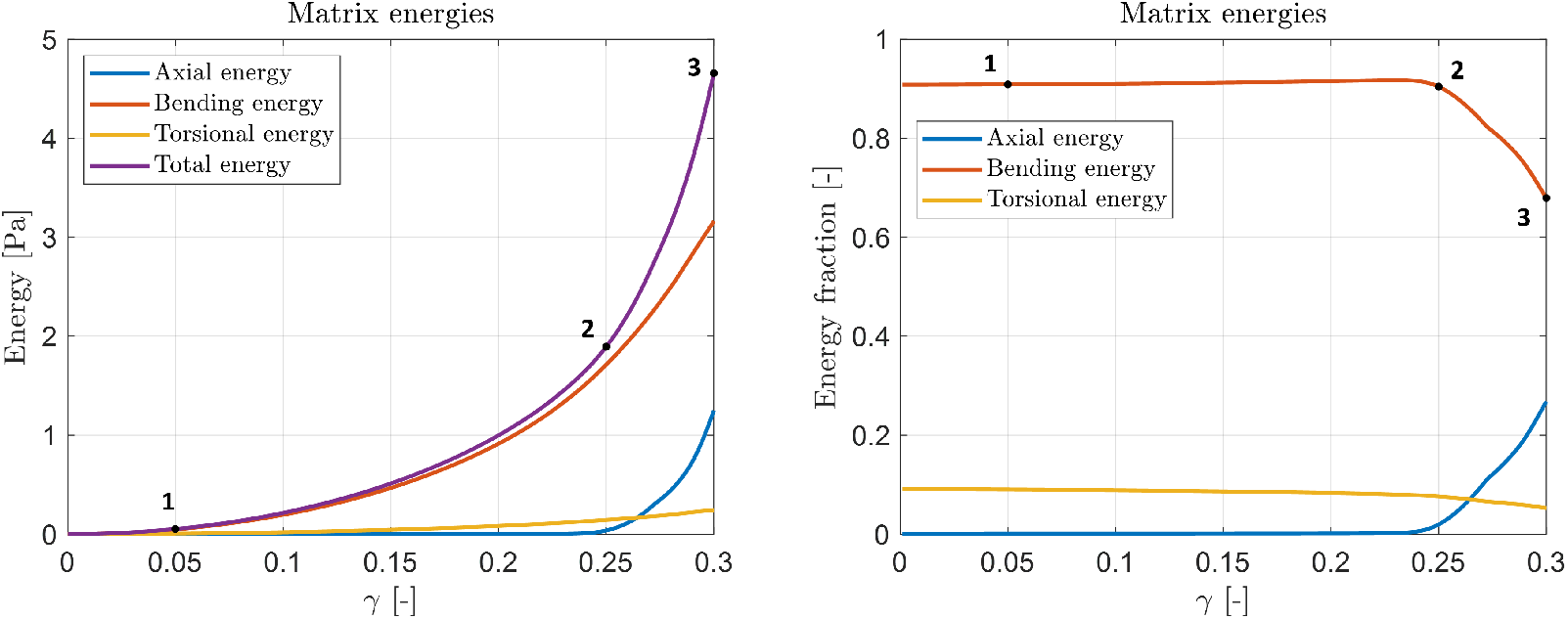
Multiscale energies: Macroscopic. Total, axial, bending and torsional accumulated strain density energies developed by the microstructure for simple shear test of the baseline matrix.

On the other hand, Figure 12 shows the accumulated multiscale energies in the fibers of the microstructure of the matrix for different shear deformation points, as defined in Figure 11. The shear behavior of the baseline matrix, the evolution of its microstructure, and both the axial, bending and torsional accumulated energies in the fibers; can be seen in videos S5, S6 and S7 of the supplementary material. It can be seen in Figure 11 that bending energy dominates in all the analyzed deformation regime, except the strain stiffening region. As in the tensile test, this energy in employed to fiber unrolling and fiber alignment with load, as corroborated in the microstructure evolution by Figure 12 and video S6. Again, the torsional energy is almost negligible in all the analyzed deformation range, according to Figures 11 and 12.

**Figure 12.**
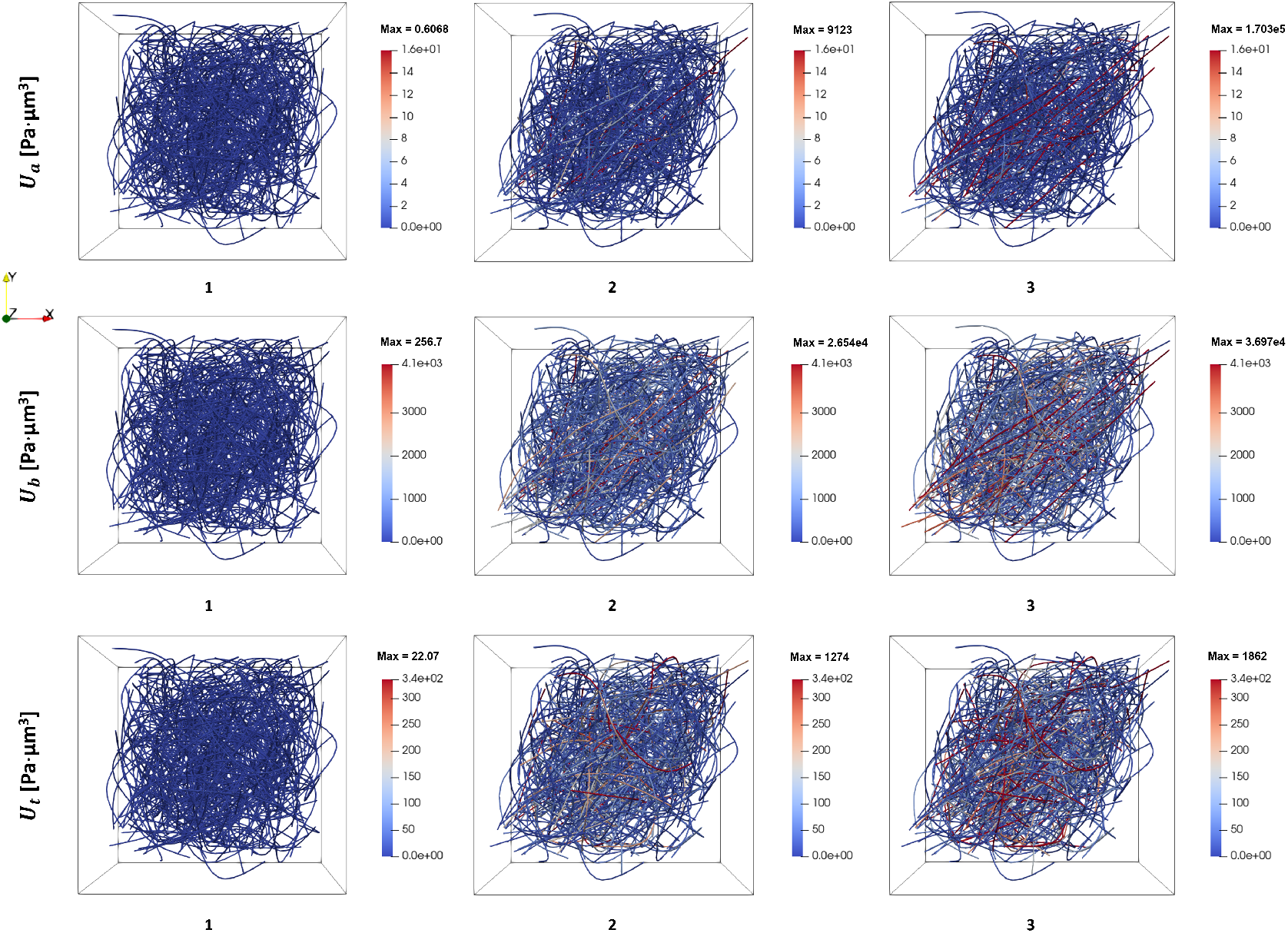
Multiscale energies: Microscopic (fiber scale). Plot in the microstructure of the axial (first row), bending (second row) and torsional (third row) of the accumulated energies in the fibers of the baseline matrix for simple shear test. Results are shown at different deformation points (1,2,3) defined in Figure 11.

It can be seen in Figures 8, 11 that strain density axial energy start being important for a certain deformation level in which fibers become straight (see Figures 9 and 12). Indeed, that the main contribution to axial energy is due to few fibers that become unrolled and straight at the end of the analysis. On the contrary, strain density bending energy is progressively increased by the contribution of fibers as they are unrolling, being of course the highest values for unrolled/straight fibers (see Figures 9 and 12). However, despite many more fibers contribute to bending than axial energies at the end of the analysis (strain stiffening region), both energies are comparable as the axial stiffness of the fibers is highest than the bending one.

### 3.5. Analysis of matrix anisotropy

Fibered matrices generated in previous sections considered the matrix generator exposed in section 3.1. This matrix generator assumes randomly oriented isolated fibers (see Box 1), so the generated matrices may be assumed as isotropic. In this section, we implement a modification of the matrix generator by enforcing the elevation angle (orientation) versus the vertical axis (being 0^*o*^ the vertical axis) of isolated fibers, to follow a normal distribution with zero mean and varying standard deviation. Therefore, the vertical axis is a preferential axis for the isolated fibers of the matrix. Figure 13 shows distribution of the elevation angle (orientation) of matrix isolated fibers, as well as their corresponding microstructures. These anisotropic microstructures were genereted with baseline matrix parameters as in previous sections, and are used in this section to analyze the effect of a preferential orientation of fibers within the matrix, i.e. anisotropy, in the mechanical behavior of matrices at tensile/compression and shear tests.

**Figure 13.**
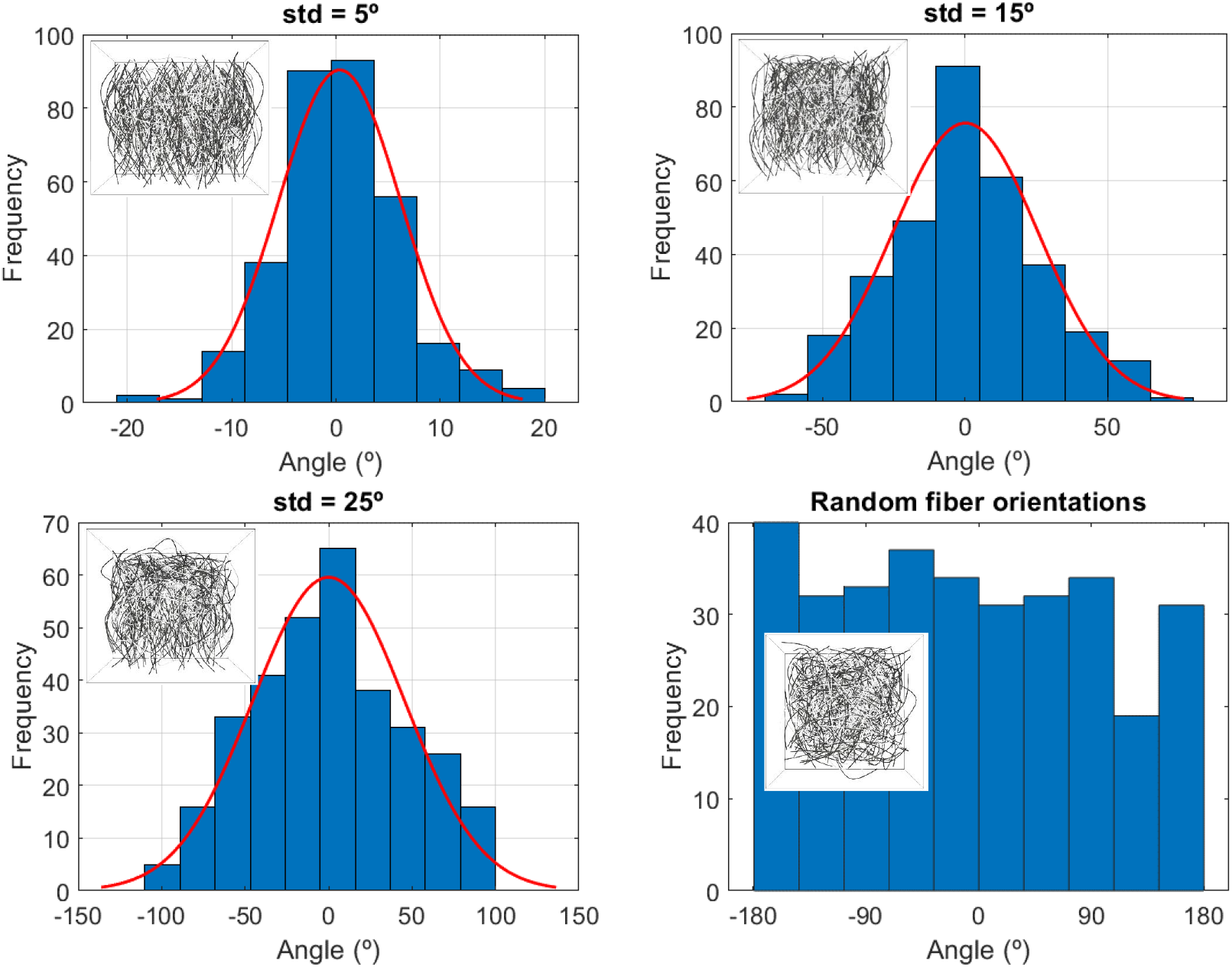
Distribution of the elevation angle (orientation) versus the vertical axis (being 0^*o*^ the vertical axis) of matrix isolated fibers, both for random distribution (isotropic matrices) and following a normal distribution for several values of the standard deviation (std). Insets in the figures are referred to the obtained matrix microstructures with baseline parameters. Dark fibers in the microstructure represents isolated fibers, whereas white fibers represent crosslink fibers.

Figure 14a shows tensile/compression tests results for different standard deviation angles of the isolated fibers of the matrices. In this case, load direction is aligned with fibers preferential direction (vertical axis). Generically, it can be seen that matrices are stiffer in all regions (compression, linear tension and strain stiffening) as fibers are more aligned with the preferential axis (lower values of the standard deviation).

**Figure 14.**
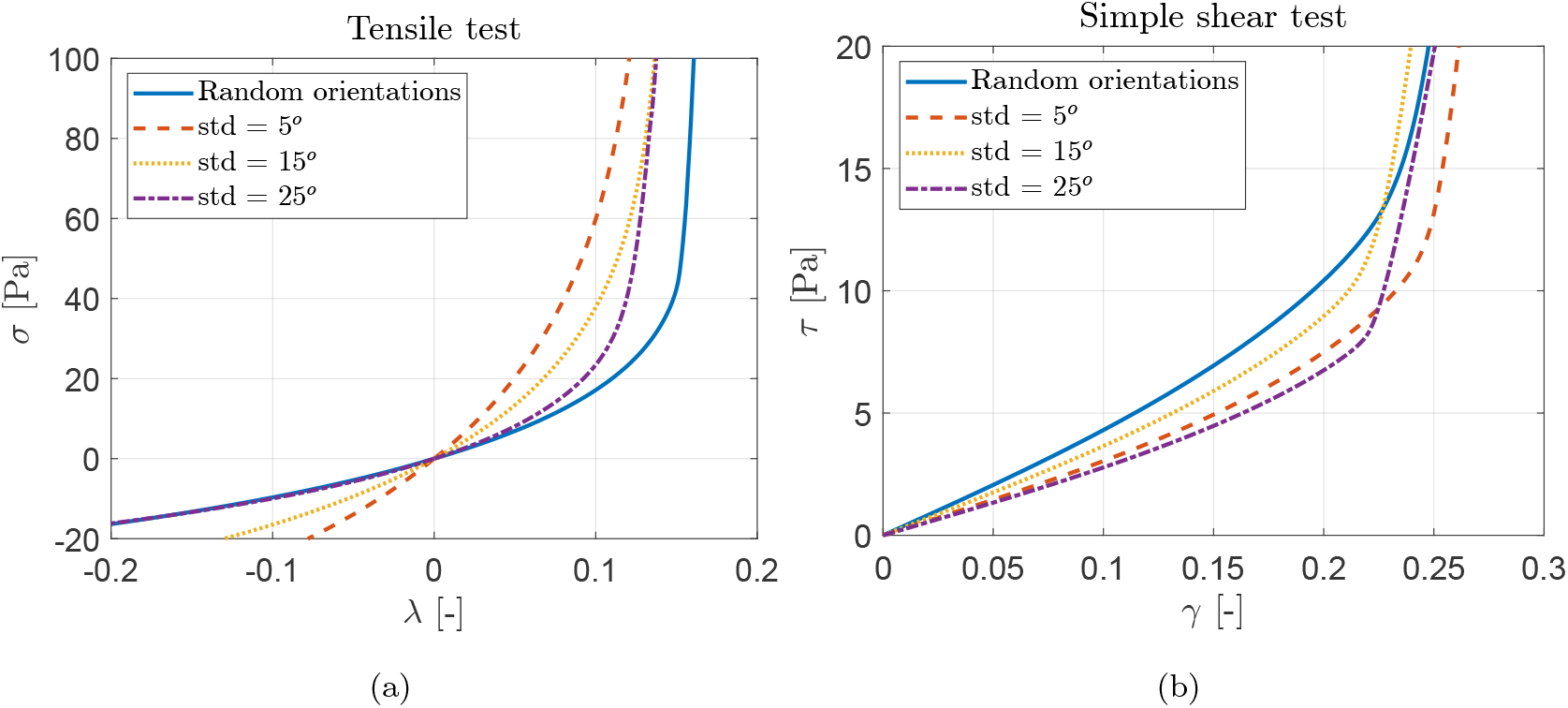
Preferentially fibered-oriented matrices (anisotropy) tests: (a) baseline matrix at tensile/compression tests for different standard deviation angles (normal distribution) of the isolated fibers along the vertical axis (see Figure 13). (b) baseline matrix at simple shear tests for different standard deviation angles (normal distribution) of the isolated fibers along the vertical axis (see Figure 13).

On the other hand, Figure 14b shows simple shear tests results for different standard deviation angles of the isolated fibers of the matrices. Load direction is parallel to the horizontal axis, whereas fibers are preferentially aligned with the vertical axis. In this case, we don’t observe a clear trend of matrices stiffening in relation to the alignment of fibers with the preferential axis.

## 4. Validation

### 4.1. Single fiber model validation

In this section the single fiber model, developed in section 2, is validated with results obtained from the literature. A first validation includes the experimental results available in single collagen fibers in chordæ tendineæ [10]. Figure 15a shows the comparison of the mechanical behavior obtained by the model versus the experimental results, for an axially stretched single fiber. Model microstructural parameters were set to *d* = 70 nm and *L* = 10 *μ*m for porcine mitral-valve chordæ tendineæ, according to the literature [49]. Fiber elastic modulus (*E* = 33 MPa) was taken from Ref. [10]. Finally, the model parameter *L*_*f*_ */L* was considered a fitting parameter, with a fitted value of 1.125, as it was difficult to find this information in the literature. As described in the parametrical analysis, *L*_*f*_ */L* determines the stretch onset in which the fiber is straight (unrolled) and the deformation of the is fiber exclusively axial (strain stiffening regime).

**Figure 15.**
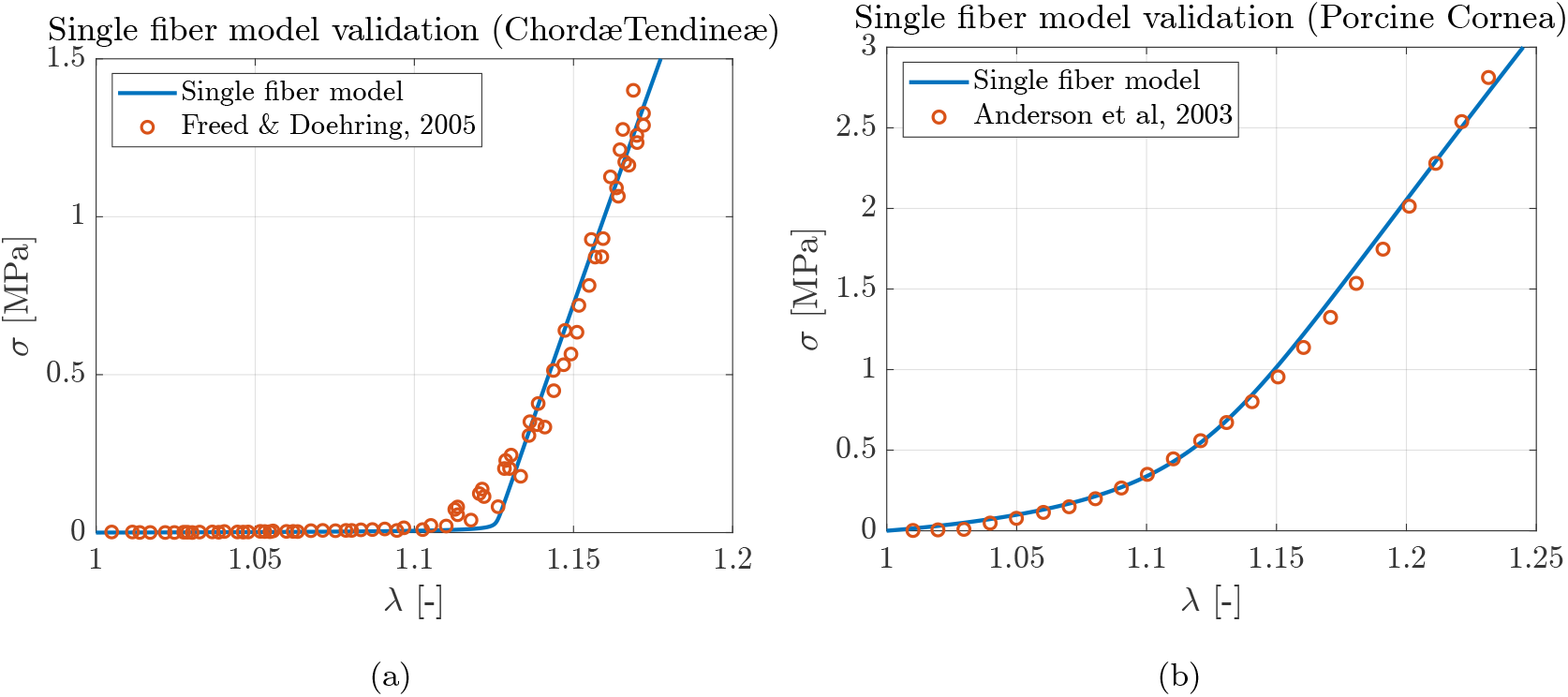
Chordæ tendineæ stress-stretch curve comparing individual randomly-curved collagen fiber model versus experimental results [10]. (b) Porcine cornea stress-stretch curve comparing individual randomly-curved collagen fiber model versus experimental results [50].

In a second validation, porcine cornea collagen fiber stress-stretch experimental results [50] were reproduced in Figure 15b. In the model simulation, the considered porcine cornea fiber diameter was *d* = 33 nm [51], whereas the elastic modulus was *E* = 27 MPa [11]. Due to lack of availability of data regarding porcine cornea fiber length, a length of *L* = 0.561 *μm* was estimated, to keep within the order of magnitude of the fiber length/diameter aspect ratio reported in the literature [52, 53]. Again, *L*_*f*_ */L* was considered a fitting parameter and set 1.11.

A good fitting of the single fiber model versus the considered experimental results can be observed in Figure 15.

### 4.2. Fibered matrix model validation

The fibered matrix model, developed in section 3, is qualitatively validated in this section with results obtained in the literature. In particular, the experimental tensile test for collagen I extracellular matrices shown in Roeder et al. [4] was used in our validation. Figure 16 shows the comparison of the mechanical behavior obtained by the fibered matrix model versus the experimental results, for an axially stretched (tensile) extracellular matrix. Model parameters considered in the simulations were set to *L* = 10 *μm, d* = 0.7 *μm, E* = 28.3 MPa, *L*_*f*_ */L* = 1.10, in order to keep within the order of magnitude and values fitted and validated in the previous section for single fibers. The fiber density of the matrix was estimated to *V*_*c*_ = 2.47 · 10^6^ fibers*/mm*^3^, due to the lack of information of this parameter in the referred experimental tests. A random distribution of fibers, without any preferential direction, was assumed in the simulations.

**Figure 16.**
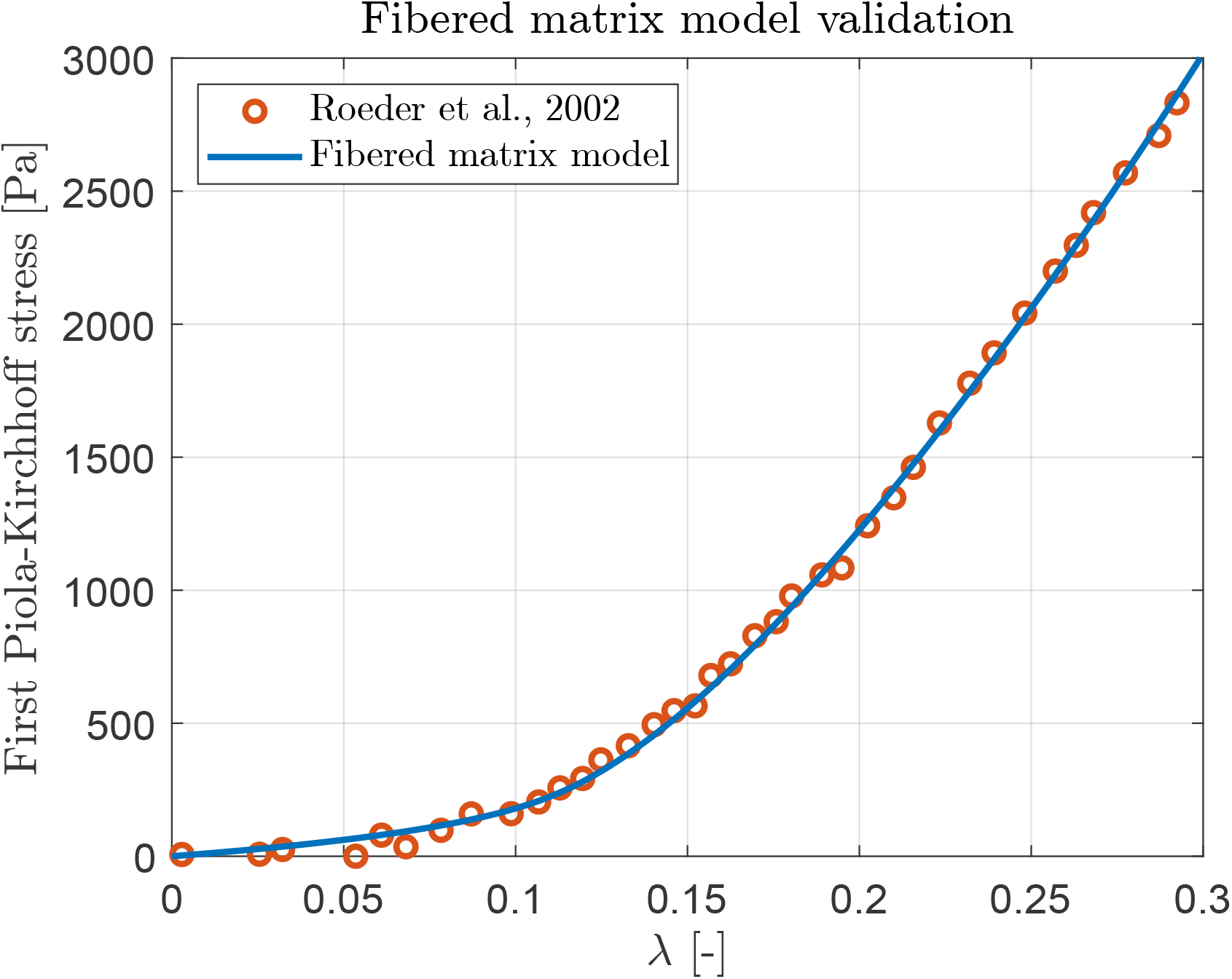
Stress-stretch tensile test comparing fibered matrix model versus collagen I extracellular matrix experimental results [4].

Again, an excellent fitting of the fibered matrix model versus the experimental results can be observed in Figure 16.

## 5. Discussion and conclusions

In this paper, a multiscale approach available to obtain the mechanical behavior of curved fibered structures was shown. The study underlines the importance of considering the curvature of fibers in order to naturally recover instabilities at compression (buckling), as well as the implications of fiber curvature in the development of strain stiffening regions at tensile and simple shear tests. Indeed, the bending energy associated to fiber unrolling, is the most important source of energy developed by fibers for the analyzed cases in tensile and shear in all deformation regions (except the strain stiffening region), whereas bending energy dominates at compression too during buckling. The numerical framework was established under an updated lagrangian framework, so key physiological processes that take place in fibered biological matrices, such as remodeling, can be easily incorporated along the formulation.

The cases of study and applications shown in the paper were primarily referred to collagen ECMs in which the domain is composed, mainly, by curved fibers and aqueous medium (not considered in the modeling due to its negligible stiffness). However, other biphasic biological matrices, coming from tissues in which the contribution of the base matrix that fibers are embedded cannot be neglected, can be straightforwardly considered in our approach. In this case, under the assumption of homogeneous base matrix and affine motion between base matrix and fibers, the macroscopic stress tensor that characterizes the mechanical behavior of the matrix for a certain macroscopic deformation state, is just added to the multiscale obtained stress tensor of the fibered domain. Moreover, this mentioned approach for biphasic fibered structures can be directly translated to study the mechanical behavior in applications outside of the biological range.

The effect of fiber alignment with the preferential direction, which is widely present in biological structures, was analyzed in this work. However, fiber viscoelasticity, which is also present in fibers, has not been considered in our study. Our future work includes the extension to fiber viscolasticity, by implementing the viscoelastic component in the nonlinear rod elements of the fibers in our formulation. Moreover, matrix viscoelasticity, which is also relevant in biology, can be accounted for in our approach just by selecting a macroscopic viscoelastic model for the mechanical behavior of the base matrix (under the assumptions mentioned above).

A parametric analysis was established both to single fiber and fibered matrices at tensile/compression and shear tests, in order to evaluate the impact of model parameters in their mechanical behavior. It can be seen that a wide spectrum of mechanical behaviors of both single fibers and fibered matrices can be reproduced, just by selecting appropriate model parameters. In addition, a qualitative validation was performed, both for single fiber and fibered matrices, showing a good agreement of the model versus experiments for available data taken from the literature.

All model parameters included in the presented approach are measurable quantities with a physical meaning. Although model parameters were selected in our analysis within the range of collagen matrices, they can be directly obtained from image segmentations and used to generate the matrix microstructure with the proposed matrix generator. Alternatively, fibered matrix microstructure may be ideally reconstructed for ECMs using advanced microscopy techniques, such as second harmonic generation (SHG) microscopy or focus ion beam scanning electron microscopy (FIB-SEM).

In summary, the proposed approach may be a useful tool to perform virtual tests to obtain the mechanical behavior of curved fibered structures present in biology and engineered materials, as well as to be implemented in multiscale FE^2^ simulations in these applications.

## Supporting information

Captions for the supplementary videos

## Acknowledgements

A.A.-F. and J.A.S.-H. gratefully acknowledge the Financial support from project PID2021-126051OB-C42 by the Ministerio de Ciencia e Innovación (MCI), Agencia Estatal de Investigación (AEI) and Fondo Europeo de Desarrollo Regional (FEDER); and project P20-01195 funded by the Consejería de Economía, Conocimiento, Empresas y Universidad de la Junta de Andalucía. E.R.-R. gratefully acknowledge the Financial support from project PID2020-113790RB-I00 by the Ministerio de Ciencia e Innovación (MCI) and Agencia Estatal de Investigación (AEI).

## Notes

### Competing Interest Statement

The authors have declared no competing interest.

