## Supplementary material for "Multiscale characterization of the mechanics of curved fibered structures with application to biological materials": Captions for the supplementary videos

**SUPPLEMENTARY INFORMATION – VIDEO CAPTIONS**

**Supplementary video S1.** Tensile/compression test of elastic single randomly curved fiber (baseline case), with parameters L = 20 µm, d = 0.2 µm, E = 100 MPa and Lf / L = 1.1.

**Supplementary video S2.** Tensile/compression tests of elastic randomly curved fiber matrices, for the baseline case with parameters L = 30 µm, d = 0.4 µm, E = 150 MPa, and Lf / L = 1.15 and $V_c$ = $2.05 \cdot 10^6$ fibers/$mm^3$. The video shows the evolution of the microstructure, as well as the axial energy accumulation in the fibers along the tests.

**Supplementary video S3.** Tensile/compression tests of elastic randomly curved fiber matrices, for the baseline case with parameters L = 30 µm, d = 0.4 µm, E = 150 MPa, and Lf / L = 1.15 and $V_c$ = $2.05 \cdot 10^6$ fibers/$mm^3$. The video shows the evolution of the microstructure, as well as the bending energy accumulation in the fibers along the test.

**Supplementary video S4.** Tensile/compression tests of elastic randomly curved fiber matrices, for the baseline case with parameters L = 30 µm, d = 0.4 µm, E = 150 MPa, and Lf / L = 1.15 and $V_c$ = $2.05 \cdot 10^6$ fibers/$mm^3$. The video shows the evolution of the microstructure, as well as the torsional energy accumulation in the fibers along the test.

**Supplementary video S5.** Simple shear test of elastic randomly curved fiber matrices, for the baseline case with parameters L = 30 µm, d = 0.4 µm, E = 150 MPa, and Lf / L = 1.15 and $V_c$ = $2.05 \cdot 10^6$ fibers/$mm^3$. The video shows the evolution of the microstructure, as well as the axial energy accumulation in the fibers along the tests.

**Supplementary video S6.** Simple shear test of elastic randomly curved fiber matrices, for the baseline case with parameters L = 30 µm, d = 0.4 µm, E = 150 MPa, and Lf / L = 1.15 and $V_c$ = $2.05 \cdot 10^6$ fibers/$mm^3$. The video shows the evolution of the microstructure, as well as the bending energy accumulation in the fibers along the tests.

**Supplementary video S7.** Simple shear test of elastic randomly curved fiber matrices, for the baseline case with parameters L = 30 µm, d = 0.4 µm, E = 150 MPa, and Lf / L = 1.15 and $V_c$ = $2.05 \cdot 10^6$ fibers/$mm^3$. The video shows the evolution of the microstructure, as well as the torsional energy accumulation in the fibers along the test.
